# Multivariate Characterization of Morpho-biometric Traits of Indigenous Helmeted Guinea Fowl (*Numida meleagris*) in Nigeria

**DOI:** 10.1101/2021.11.24.469826

**Authors:** Abdulmojeed Yakubu, Praise Jegede, Mathew Wheto, Ayoola J. Shoyombo, Ayotunde O. Adebambo, Mustapha A. Popoola, Osamede H. Osaiyuwu, Olurotimi A. Olafadehan, Olayinka O. Alabi, Comfort I. Ukim, Samuel T. Vincent, Harirat L. Mundi, Adeniyi Olayanju, Olufunmilayo A. Adebambo

## Abstract

This study was embarked upon to characterise phenotypically helmeted guinea fowls in three agro-ecologies in Nigeria using multivariate approach. Eighteen biometric characters, four morphological indices and eleven qualitative (phaneroptic) traits were investigated in a total of 569 adult birds (158 males and 411 females). Descriptive statistics, non-parametric Kruskal–Wallis H test followed by the Mann–Whitney U test for post hoc, Multiple Correspondence Analysis (MCA), General Linear Model, Canonical Discriminant Analysis, Categorical Principal Component Analysis and Decision Trees were employed to discern the effects of agro-ecological zone and sex on the morphostructural parameters. Agro-ecology had significant effect (P<0.05; P <0.01) on all the colour traits. In general, the most frequently observed colour phenotype of guinea fowl had pearl plumage colour (54.0%), pale red skin colour (94.2%), black shank colour (68.7%), brown eye colour (49.7%), white earlobe colour (54.8%) and brown helmet colour (72.6%). The frequencies of helmet shape and wattle size were significantly influenced (P <0.01) by agro-ecology and sex. Overall, birds from the Southern Guinea Savanna zone had significantly higher values (P <0.05) for most biometric traits compared to their Sudano-Sahelian and Tropical Rainforest counterparts. They were also more compact (120.83±1.61 vs. 113.96±0.97 vs. 111.33±1.19) and had lesser condition index (8.542±0.17 vs. 9.92±0.10 vs. 9.61±0.13) than their counterparts in the two other zones. The interaction between agro-ecology and sex had significant effect (P <0.05) on some quantitative variables. The MCA and discriminant analysis revealed considerable intermingling of the phaneroptic, biometric traits and body indices especially between the Sudano-Sahelian and Tropical Rainforest birds. Inspite of the high level of genetic admixture, the guinea fowl populations could best be distinguished using wing length, body length and eye colour. However, further complementary work on genomics will guide future selection and breeding programmes geared towards improving the productivity, survival and environmental adaptation of indigenous helmeted guinea fowls in the tropics.

## Introduction

Poultry species serve as important sources of animal protein and household income, especially for low-input and marginalized rural communities [1,2,3]. The helmeted guinea fowl (*Numida meleagris*) belongs to the Galliformes order and the Numididae family. The game bird is terrestrial and commonly found in Africa [4]. The birds are indigenous to West Africa North of the Equatorial forest and are believed to have originated from the coast of Guinea in West Africa [5]. Based on evidence from archaeozoology and art, it was suggested that Mali and Sudan were centres of domestication of this species and this might have occurred about 2,000 years BP [6]. In Nigeria, the Guinea fowl is a common game bird found mainly in the Savanna region of Northern Nigeria [7]. Guinea fowl farmers are basically involved in three major production systems: These include the Extensive System (Free range), Semi-intensive System (Partial confinement) and the Intensive System (Complete enclosure) [8]. In comparison with chicken, guinea fowl is economically more attractive in the tropics because it is not very demanding in terms of its diet, more rustic and adapts better to traditional farming system [9,10,11]. Guinea fowl is also highly valued for its meat and eggs. The meat is rich in vitamins and contains less cholesterol and fats, thereby making it a high quality protein source [12]. Additionally, the bird is used for different cultural purposes, and plays a role in poverty reduction among rural dwellers [13]. The bird also breeds seasonally and reaches its peak breeding activity during the summer period [14].

Every livestock species or breed is a veritable component of the animal genetic diversity of the world that deserves immense attention [15]. Despite the usefulness of guinea fowl, it is poorly characterised in the tropics. This has limited its value as an unexploited potential for economic and industrial growth. Therefore, there is need for proper characterisation geared mainly towards improvement in meat and egg production. The first step in such characterisation as outlined by FAO [16] involves the use of phenotypic characteristics which are aspects of physical appearance or other body parameters that can be measured qualitatively and quantitatively [17]. Variations in phenotypes have remained [18], and tolerance or susceptibility of birds to stressful environment could be linked to their phenotypic traits [19, 20]. Hence, the need to understand such phenotypic diversity in the helmeted Guinea fowls especially in populations that have adapted to local environmental conditions. Under resource-poor settings, phenotypic approach is fundamental in livestock management because it is simple, fast, and cost-effective [21]. Also, morpho-biometrical characterization (qualitative and quantitative traits) will enable proper selection of elite animals, breeding, conservation and sustainable use of indigenous animal resources [22, 23]. Qualitative traits such as plumage colour, skin colour, shank colour, eye colour, helmet shape, wattle possession and skeleton structure are useful to farmers and breeders for identification and classification of guinea fowl and to meet consumer preferences for specific phenotypic traits [24]. On the other hand, biometric measurements such as body weight, body length, chest circumference, wing length, wingspan and shank length are useful in breeding programmes, to revaluate local breeds, allow the preservation of animal biodiversity and support consumer demands [25, 26]. When such morphometric traits are considered jointly, multifactorial analyses have been shown to assess better the within-population variation which can be utilized in the discrimination of different population types [25, 27]. In Nigeria, south Saharan Africa, there is dearth of information on the phenotypic diversity of guinea fowls [28]. The current study, therefore, was embarked upon to characterise morphologically guinea fowl in Nigeria using qualitative traits and linear body measurements. The knowledge of the morpho-biometrical traits will support the implementation of breeding and conservation strategies in order to guarantee the survival and continuous production of the guinea fowl genetic resource in the tropics for improved food security and livelihoods.

## Materials and Methods

### Ethics Statement

In order to properly carry out the research, we adhered strictly to the ethical guidelines of the Global code of conduct for research in resource- poor settings [29] following the Convention on Biological Diversity and Declaration of Helsinki. Although the study did not involve the collection of blood and other tissue samples, we obtained field approval from the Research and Publication Directorate of Nasarawa State University, Keffi through permit no NSUK/FAC/ANS/GF100.

### Study Area

The study was carried out in Nasarawa State and Abuja (Southern Guinea Savanna zone), Bauchi and Kano States (Sudano-Sahelian zone) and Ogun and Oyo States (Tropical Rainforest zone) of Nigeria. The choice of sites was informed by the relative availability of indigenous and exotic guinea fowls and ease of data collection. The climates and vegetations of Nasarawa State and Abuja have been described in an earlier study [30]. Bauchi State is located strategically between latitudes 9°30’ and 12°30’ North of the equator and between longitudes 8°45’ and 11°0’ East of the Greenwich meridian. The State has two distinct ecological zones; namely, the Sudan savannah and the Sahel savannah. While the Sudan savannah covers the Southern part of the State where the vegetation gets richer and richer towards the South, the Sahel (semi-desert vegetation) covers the Western and Northern parts of the State, with characteristic features of isolated strands of horny shrubs and sandy soils [31]. The mean daily temperature is highest in April (30.0°C) and lowest in December (22.1°C) [32]. The average relative humidity ranges from 35% in February to 94% in August. Monthly rainfall ranges from 0.0mm in December and 0.5mm in January to about 340mm in August. Kano State is part of the Sudano-Sahelian zone of Nigeria and is located approximately between longitudes 8° 45 E and 12° 05 E latitudes 10° 30 N and 13° 02 N. The natural vegetation is a mixture of Sudan Savannah and Sahel thorn shrubs species, which are sparsely distributed over the entire area, with variation in density from one place to another [33]. The mean annual rain-fall is within the range of 800 mm to 900 mm, with the wet season lasting from May to mid-October and peaks in August while the dry season extends from mid-October to mid-May. The mean annual temperature is about 26°C [34].

Ogun State located between on latitude 6°12′N and 7°47′N and longitude 3°0′E and 5°0′E. Vegetationally, the State is characterized mainly by tropical rainforest,. The average temperature value varies from one month to another, with a minimum average of 25.7 °C in July and a maximum of 30.2 °C in February. The State has two distinct seasons (wet and dry) with mean annual rainfall value ranging between 1,400 and 1,500 mm [35]. Oyo State is found on latitude 8°.00’N and longitude 4°.00’E. It is characterized by humid rain forest vegetation. The average temperature ranges between 25.0 °C and 35.0 °C. The wet season starts from April and ends in October while the dry season lasts from November to March. The rainfall pattern is bimodal with annual mean value of 1250 mm [36].

### Sampling Procedure

A total of five hundred and sixty nine (569) adult (8 months of age) Nigerian indigenous guinea fowls [Southern Guinea Savanna: 109 birds (27 males and 82 females); Sudano-Sahelian: 290 birds (80 males and 190 females) and Tropical Rainforest: 190 birds (51 males and 139 females)] were utilized in the study. The indigenous birds were randomly sampled in smallholder rural farmers flocks and managed under the traditional low-input settings. Multistage sampling procedure was purposively and randomly adopted in the selection of States, Local Government Areas (LGAs), villages and guinea fowl keepers in each agro-ecological zone. States, LGAs and villages were purposively selected based on the knowledge of the availability of guinea fowls in the communities as provided by the local Extension Agents and Community Heads. The number of sampling locations varied [4 LGAs and 11 villages (Southern Guinea Savanna); 5 LGAs and 15 villages (Sudano-Sahelian); 4 LGAs and 13 villages (Tropical Rainforest)]. Based on willingness to participate in the research, eleven individuals were then randomly selected from each village making a total of 429 households (n=121, 165 and 143 for Southern Guinea Savanna, Sudano-Sahelian and Tropical Rainforest, respectively).

### Data Collection

Data collection was done in the rainy season month of April to June, 2020. Morphologically distinct Guinea fowls were identified using phenotypic traits based on the standard descriptors by FAO [16], AU-IBAR [37] and the colour chart of Guinea fowl by GFIA [38]. The sexes were distinguished through visualisation of the vent and the use of helmet shape as well as wattle size and shape [28]. Eleven qualitative (phaneroptic) parameters such as plumage colour, skin colour, shank colour, eye colour, earlobe colour, helmet colour, helmet shape, wattle possession, wattle size, wattle shape and skeletal structure were used to characterize the guinea fowls morphologically. For quantitative (biometric) description, the following body parts were measured:

Body weight (kg): The live weight of the guinea fowl.

Head length (cm): Taken between the most protruding point of the occipital and the frontal (lacrimal) bone.

Head thickness (cm): Head thickness measured as the circumference at the middle of the head

Helmet length (cm): Measured as the distance between the base of the head to the tip of the helmet

Helmet width (cm): Measured as the distance between the broadest part of the helmet

Wattle length (cm): Taken as the distance between the base of the beak and the tip of the wattle

Wattle width (cm): Measured as the distance between the broadest part of the wattle

Neck length (cm): Distance between the occipital condyle and the cephalic borders of the coracoids.

Neck circumference: Taken at the widest point of the neck

Wing length (cm): Taken from the shoulder joint to the extremity of the terminal phalanx, digit 111.

Wing Span: Distance between the two wings when stretched out.

Body length (cm): The distance from the first cervical vertebra (Atlas) to the posterior end of the ischium

Trunk Length (cm): The distance between shoulder joint and posterior edge of the ischium,

Keel length (cm): Keel length (Sternum or breast bone) measured from the anterior point of the keel to the posterior end.

Chest circumference (cm): This was taken as the circumference of the body around the breast region.

Thigh length (cm): Distance between the hock joint and the pelvic joint;

Thigh circumference (cm): Measured as the circumference at the widest point of the thigh;

Shank length: Shank length was measured as the distance between the foot pad and the hock joint.

Shank thickness: Shank thickness was measured as the circumference at the middle or widest part of the shank.

Also, the following conformation indices were estimated [39]:

Massiveness: The ratio of live body weight to trunk length x 100

Compactness: The ratio of chest circumference to trunk length x 100

Long-leggedness: The ratio of shank length to body length x 100

Condition index: The ratio of live body weight to wing length × 100.

The weight measurement was taken using a hanging digital scale (WeiHeng Brand), the width measurements were taken using a vernier caliper (0.01 mm precision) while the length and circumference measurements were taken using a flexible tape measure.

## Statistical Analysis

### Descriptive Statistics

Descriptive statistics were computed to determine the frequencies of the qualitative traits. Where statistical significant differences in the frequencies were obtained at agro-ecological and sex levels, they were assessed using the non-parametric Kruskal–Wallis H test followed by the Mann–Whitney U test for post hoc separation [40] of IBM-SPSS software (2020).

### Correspondence Analysis

Multiple Correspondence analysis (MCA) was used to establish the relationships between the qualitative traits: Wattle possession and skeleton structure had zero variance and were excluded from the MCA using JMP 16 [41] statistical software.

### Factorial Analysis

General linear model (GLM) of IBM-SPSS software [42] was employed to test the fixed effects of agro-ecology and sex as well as their interaction on quantitative variables. Significant means were separated using Least Significant Difference (LSD) method at P<0.05 level.

The general linear model employed was:

Y_ijk_ = μ + A_i_ + S_j_ + (AS)_ij_ + e_ijk_

Y_ijk_ = individual observation

μ = population mean

A_i_ = i^th^ agro-ecology fixed effect (i = southern guinea savanna, sudano-sahelian, tropical rainforest). S_j_ = j^th^ sex fixed effect (j = male, female)

(AS)_ij_ = i^th^ agro-ecology and j^th^ sex interaction effect

e_ijk_ = random error associated with each record

### Stepwise Canonical Discriminant Analysis

Canonical discriminant analysis [43] option of IBM-SPSS [42] statistical software was applied to classify birds in the three agro-ecological zones based on quantitative traits. In the analysis, all the eighteen biometric traits and four conformation indices (covariates) were entered in a stepwise fashion as explanatory variables to establish and outline population clusters [44] based on agro-ecology. F-to-remove statistics was the criterion for variables’ selection while multicollinearity was detected among the variables in the discriminant function using tolerance statistics. The ability of this discriminant model to identify birds in the Southern Guinea Savanna, Sudano-Sahelian and Tropical Rainforest zones was indicated as the percentage of individuals correctly classified from the sample that generated the model. The accuracy of the classification was evaluated using split-sample validation (cross-validation).

### Categorical Principal Component Analysis

Categorical principal component analysis (CATPCA) procedure was employed to explore hidden relationships among the qualitative and quantitative traits as described by Martin-Collado et al. [45]. This was to allow for appropriate grouping of the guinea fowls based on agro-ecology and sex. The PCs were extracted based on Eigen values greater than 1 criterion. The convergence was 0.00001 with maximum iterations of 100. The PC matrix was rotated using the varimax criterion with Kaiser Normalization to facilitate easy interpretation of the analysis. The reliability of the PCA was tested using Chronbach’s alpha using IBM-SPSS [42].

### Decision Trees

CHAID and Exhaustive CHAID algorithms were employed to assign the birds into agro-ecological zones using the qualitative and quantitative traits as the predictor variables. CHAID is a tree-based model with merging, partitioning and stopping stages that recursively uses multi-way splitting procedures to form homogenous subsets using Bonferroni adjustment until the least differences between the predicted and actual values in a response variable are obtained [46]. It produces terminal nodes and finds the best possible variable or factor to split the node into two child nodes. The Exhaustive CHAID, as a modification of CHAID algorithm, applies a more detailed merging and testing of predictor variables [47]. The goodness-of-fit criterion to assess the efficiency of the CHAID and Exhaustive CHAID models was the risk value including its associated standard error. IBM-SPSS [42] software was also used for the Decision Trees’ analysis

## Results

### Distribution of the Qualitative Traits

The frequency distribution of the colour traits of indigenous helmeted guinea fowl are shown in Table 1. Agro-ecology significantly affected (P<0.05; P <0.01) all the six traits investigated. No definite pattern of variation in each class of the colour traits was observed among the three agro-ecological zones. Generally, the most frequent colour phenotype of helmeted guinea fowl in Nigeria had pearl plumage colour (54.0%), pale red skin colour (94.2%), black shank colour (68.7%), brown eye colour (49.7%), white earlobe colour (54.8%) and helmet colour (72.6%). However, sex did not significantly influence (P>0.05) all the six colour traits.

**Table 1.**
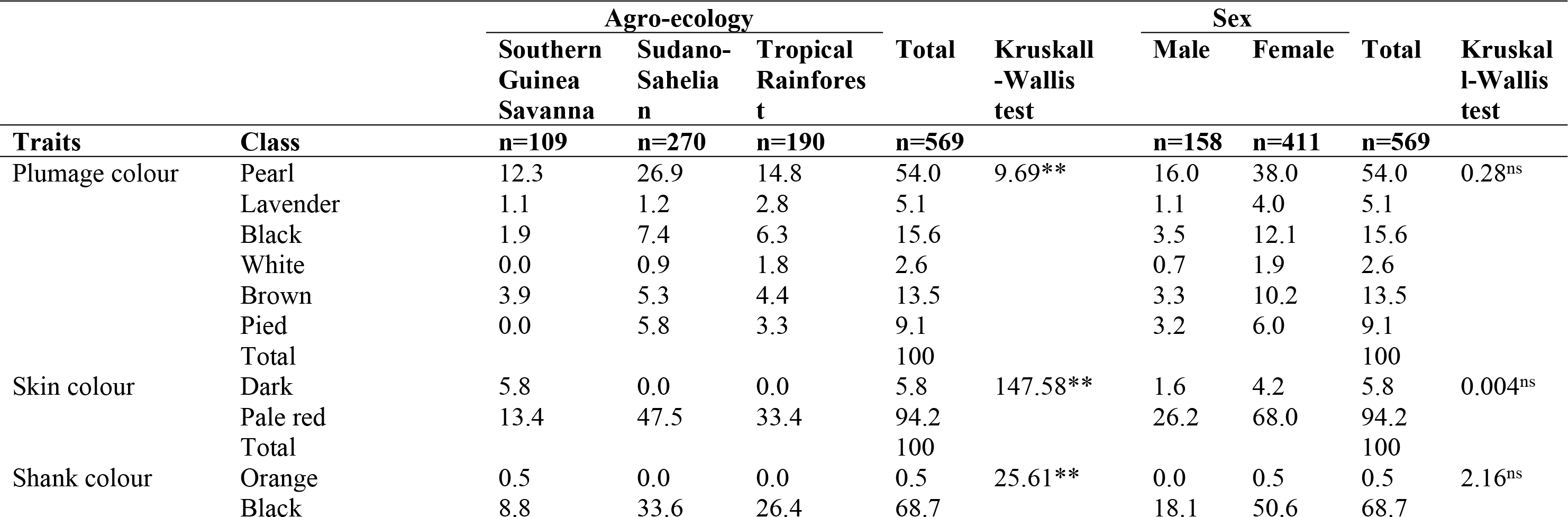
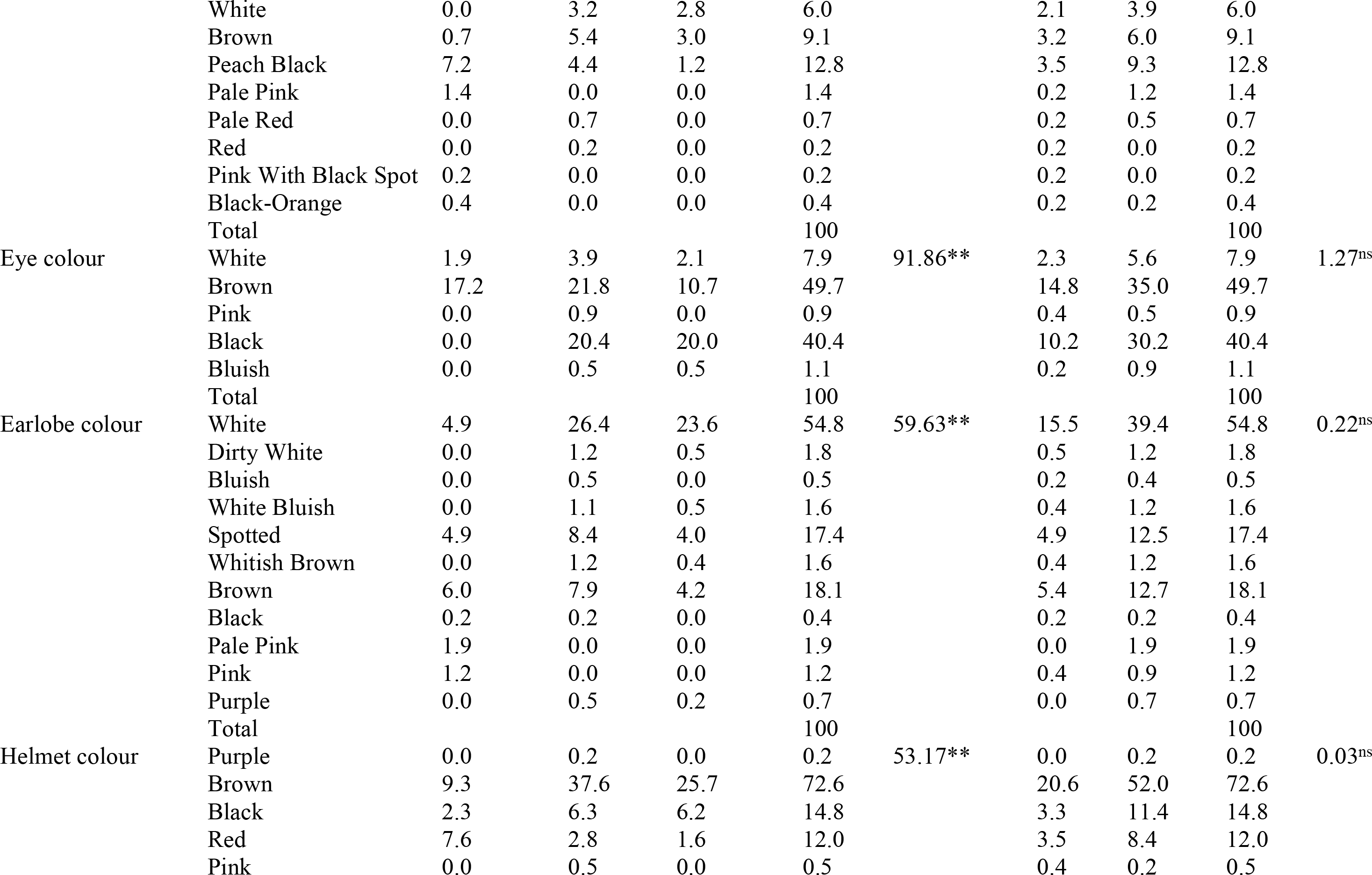
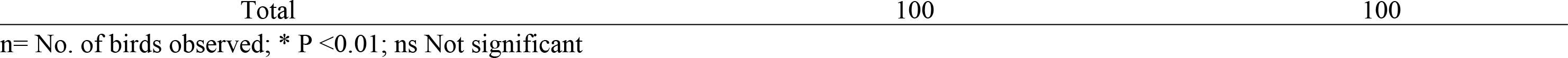
Frequency (%) of colour traits of indigenous helmeted guinea fowl based on agro-ecology and sex

The frequencies of helmet shape and wattle size were significantly affected by agro-ecology (P <0.01) (Table 2). While most of the birds had single helmet shape (50.8%), which appeared to be more in the Sudano-Sahelian and Tropical Rainforest zones, wattle size did not follow a definite pattern. All the birds in the three agro-ecologies had wattle and were skeletally normal (P >0.01). However, sex had a significant effect (P <0.01) on helmet shape (where more females were single), wattle size (where that of males appeared larger) and wattle shape (where more females carried theirs flat).

**Table 2.** Frequency (%) of helmet shape, wattle possession, size and shape including skeletal structure of indigenous helmeted guinea fowl based on agro-ecology and sex

| Traits | Class | Agro-ecology |  |  | Total | Kruskal<br>l-Wallis<br>test | Sex |  | Total | Kruskal<br>-Wallis<br>test |
| --- | --- | --- | --- | --- | --- | --- | --- | --- | --- | --- |
|  |  | Southern<br>Guinea<br>Savanna<br>n=109 | Sudano-Sahelia<br>n=270 | Tropical<br>Rainfores<br>t<br>n=190 |  |  | Male<br>n=158 | Female<br>n=411 |  |  |
| Helmet shape | Slanted Backward | 13.0 | 5.3 | 4.0 | 22.3 | 43.61** | 6.2 | 16.2 | 22.3 | 94.57** |
|  | Single | 0.2 | 29.3 | 21.3 | 50.8 |  | 1.8 | 49.0 | 50.8 |  |
|  | Erect | 6.0 | 12.8 | 8.1 | 26.9 |  | 19.9 | 7.0 | 26.9 |  |
|  | Total |  |  |  | 100 |  |  |  | 100 |  |
| Wattle possession | Present | 19.2 | 47.5 | 33.4 | 100.0 | 0.00<sup>ns</sup> | 27.8 | 72.2 | 100 | 0.00<sup>ns</sup> |
|  | Absent | 0.0 | 0.0 | 0.0 | 0.0 |  | 0.0 | 0.0 | 0.0 |  |
|  | Total |  |  |  | 100 |  |  |  | 100 |  |
| Wattle size | Large | 12.7 | 23.7 | 13.4 | 49.7 | 18.79<sup>**</sup> | 23.9 | 25.8 | 49.7 | 115.34<sup>**</sup> |
|  | Small | 6.5 | 23.7 | 20.0 | 49.7 |  | 3.9 | 46.4 | 50.3 |  |
|  | Total |  |  |  | 100 |  |  |  | 100 |  |
| Wattle shape | Cupped | 5.1 | 13.9 | 8.8 | 27.8 | 0.47<sup>ns</sup> | 27.2 | 0.5 | 27.8 | 526.89<sup>**</sup> |
|  | Flat | 14.1 | 33.0 | 24.3 | 71.4 |  | 0.5 | 70.8 | 71.4 |  |
|  | Cupped Flat | 0.0 | 0.5 | 0.4 | 0.9 |  | 0.0 | 0.9 | 0.9 |  |
|  | Total |  |  |  | 100 |  |  |  | 100 |  |
| Skeletal structure | Normal | 19.2 | 47.5 | 33.4 | 100 | 0.00<sup>ns</sup> | 27.8 | 72.2 | 100 | 0.00<sup>ns</sup> |
|  | Creeper | 0.0 | 0.0 | 0.0 | 0.0 |  | 0.0 | 0.0 | 0.0 |  |
|  | Polydactyl | 0.0 | 0.0 | 0.0 | 0.0 |  | 0.0 | 0.0 | 0.0 |  |

n= No. of birds observed; \*\* Significant at P <0.01; ns Not significant

### Biplot of the Multiple Correspondence Analysis

The MCA revealed the association between the qualitative traits and agro-ecological zones in two dimensions (Figure 1). The first dimension was high and represented 93.2% of the deviation from independence while the second dimension signified 6.8% of the total variation based on the inertia. The agro-ecological zones were not clustered perfectly (as can been in the low inertia values of 0.168 and 0.012) considering the intermingling of some qualitative traits. This was more noticeable between birds in the Sudano-Sahelian and Tropical Rainforest zones. Therefore, discrimination of the traits appears very weak. However, on the right hand side of the biplot, peach black, orange and pale pink shank colour, dark skin colour, red and slanted backward helmet seemed to be more associated with the Southern Guinea Savanna zone.

**Fig 1.** A biplot showing the relationship between the qualitative traits and agro-ecological zones

#### The Fixed Effects of Quantitative Variables

The results of the univariate analysis revealed significant effect (P*<*0.05) of agro-ecology on the biometric traits and morphological indices of the guinea fowls (Table 3). Overall, birds from the Southern Guinea Savanna zone had significantly higher values (P <0.05) for most zoometrical traits compared to their Sudano-Sahelian and Tropical Rainforest counterparts. However, the former only differed (P <0.05) from the later in body weight (91.47±0.02 vs. 1.42±0.02), wattle width (1.66±0.03 vs. 1.53±0.04) and chest circumference (29.66±0.24 vs. 28.83±0.29). As regards conformation indices, Southern Guinea Savanna birds were more compact (120.83±1.61 vs. 113.96±0.97 vs. 111.33±1.19) and had lesser condition index (8.542±0.17 vs. 9.92±0.10 vs. 9.61±0.13) than those of Sudano-Sahelian and Tropical Rainforest. Sex significantly influenced (P*<*0.05) only the biometric traits as the males had higher head thickness, helmet length, wattle length, wattle width, wing length, wing span, body length, trunk length, chest circumference and thigh length (Table 4).

**Table 3.** Effect of agro-ecology on the quantitative traits (Mean±S.E.) of indigenous helmeted guinea fowls

| Traits | Agro-ecology |  |  | P-value |
| --- | --- | --- | --- | --- |
|  | Southern Guinea Savanna | Sudano-Sahelian | Tropical Rainforest |  |
| Body weight | 1.49±0.03<sup>a</sup> | 1.47±0.02<sup>a</sup> | 1.42±0.02<sup>b</sup> | 0.048 |
| Head length | 4.49±0.05<sup>b</sup> | 4.76±0.03<sup>a</sup> | 4.71±0.04<sup>a</sup> | 0.001 |
| Head thickness | 10.84±0.07<sup>b</sup> | 11.09±0.04<sup>a</sup> | 11.04±0.05<sup>a</sup> | 0.009 |
| Helmet length | 2.31±0.06<sup>a</sup> | 1.96±0.04<sup>b</sup> | 1.99±0.05<sup>b</sup> | 0.001 |
| Helmet width | 1.64±0.04<sup>a</sup> | 1.37±0.03<sup>b</sup> | 1.33±0.03<sup>b</sup> | 0.001 |
| Wattle length | 2.36±0.05 | 2.33±0.03 | 2.30±0.04 | 0.695 |
| Wattle width | 1.88±0.06<sup>a</sup> | 1.66±0.03<sup>b</sup> | 1.53±0.04<sup>c</sup> | 0.001 |
| Neck length | 14.03±0.20<sup>a</sup> | 11.58±0.12<sup>b</sup> | 11.24±0.15<sup>b</sup> | 0.001 |
| Neck circumference | 8.02±0.13<sup>a</sup> | 7.215±0.08<sup>b</sup> | 7.226±0.09<sup>b</sup> | 0.001 |
| Wing length | 17.88±0.18<sup>a</sup> | 14.86±0.11<sup>b</sup> | 14.73±0.13<sup>b</sup> | 0.001 |
| Wing Span | 38.88±0.29<sup>a</sup> | 35.48±0.17<sup>b</sup> | 35.09±0.21<sup>b</sup> | 0.001 |
| Body length | 38.65±0.51<sup>a</sup> | 36.36±0.30<sup>b</sup> | 35.65±0.37<sup>b</sup> | 0.001 |
| Trunk Length | 26.81±0.23<sup>a</sup> | 26.14±0.14<sup>b</sup> | 25.98±0.17<sup>b</sup> | 0.010 |
| Keel length | 11.23±0.11<sup>a</sup> | 11.03±0.06<sup>ab</sup> | 10.83±0.08<sup>b</sup> | 0.008 |
| Chest circumference | 32.24±0.40<sup>a</sup> | 29.66±0.24<sup>b</sup> | 28.83±0.29<sup>c</sup> | 0.001 |
| Thigh length | 11.64±0.12<sup>a</sup> | 10.85±0.07<sup>b</sup> | 10.68±0.09<sup>b</sup> | 0.001 |
| Shank length | 7.39±0.06<sup>a</sup> | 7.04±0.03<sup>b</sup> | 7.01±0.04<sup>b</sup> | 0.001 |
| Shank thickness | 4.10±0.10<sup>b</sup> | 5.45±0.06<sup>a</sup> | 5.39±0.08<sup>a</sup> | 0.001 |
| Massiveness | 5.57±0.12 | 5.68±0.07 | 5.49±0.09 | 0.225 |
| Compactness | 120.83±1.61<sup>a</sup> | 113.96±0.97<sup>b</sup> | 111.33±1.19<sup>b</sup> | 0.001 |
| Long-leggedness | 19.17±0.23<sup>b</sup> | 19.62±0.14<sup>ab</sup> | 19.88±0.17<sup>a</sup> | 0.042 |
| Condition index | 8.542±0.17<sup>b</sup> | 9.92±0.10<sup>a</sup> | 9.61±0.13<sup>a</sup> | 0.001 |

Means within rows with P <0.05 are significantly different

**Table 4.** Effect of sex on the quantitative traits (Mean±S.E.) of indigenous helmeted guinea fowls

| Traits | Sex |  | P-value |
| --- | --- | --- | --- |
|  | Male | Female |  |
| Body weight | 1.48±0.02 | 1.44±0.01 | 0.136 |
| Head length | 4.67±0.04 | 4.64±0.02 | 0.617 |
| Head thickness | 11.08±0.05 | 10.91±0.03 | 0.006 |
| Helmet length | 2.17±0.05 | 2.00±0.03 | 0.003 |
| Helmet width | 1.48±0.03 | 1.42±0.02 | 0.131 |
| Wattle length | 2.38±0.04 | 2.28±0.03 | 0.036 |
| Wattle width | 1.77±0.04 | 1.61±0.03 | 0.003 |
| Neck length | 12.37±0.16 | 12.19±0.10 | 0.352 |
| Neck circumference | 7.57±0.10 | 7.41±0.06 | 0.151 |
| Wing length | 16.08±0.14 | 15.57±0.09 | 0.002 |
| Wing Span | 36.80±0.23 | 36.17±0.14 | 0.018 |
| Body length | 37.47±0.40 | 36.30±0.24 | 0.012 |
| Trunk Length | 26.67±0.18 | 25.95±0.11 | 0.001 |
| Keel length | 11.08±0.08 | 10.98±0.05 | 0.304 |
| Chest circumference | 30.66±0.32 | 29.82±0.19 | 0.022 |
| Thigh length | 11.21±0.10 | 10.90±0.06 | 0.006 |
| Shank length | 7.18±0.04 | 7.11±0.03 | 0.131 |
| Shank thickness | 5.06±0.08 | 4.90±0.05 | 0.098 |
| Massiveness | 5.59±0.09 | 5.57±0.06 | 0.903 |
| Compactness | 115.48±1.27 | 115.26±0.76 | 0.881 |
| Long-leggedness | 19.36±0.18 | 19.75±0.11 | 0.060 |
| Condition index | 9.33±0.14 | 9.38±0.08 | 0.754 |

Means within rows with P <0.05 are significantly different

### The Interaction Effects of Quantitative Variables

The interaction between agro-ecology and sex only had significant effect (P <0.05) on some quantitative variables where the male had an edge over the female in the three agro-ecological zones (Table 5). In the Sudano-Sahelian zone, the body weight of males (1.53±0.03) was higher than that of the females (1.42±0.02) likewise keel length (11.20±0.11 vs. 10.85±0.07), massiveness (5.86±0.12 vs. 5.51±0.08) and condition index (10.14±0.18 vs. 9.69±0.11). Similar pattern was observed in helmet length, (2.57±0.11 vs. 2.04±0.06), wattle length (2.53±0.09 vs. 2.19±0.05) and neck circumference (8.29±0.23 vs. 7.76±0.13) in the Southern Guinea Savanna zone.

**Table 5.** Agro-ecology and sex interaction effect on the quantitative traits (Mean±S.E.) of indigenous helmeted guinea fowls

|  | Southern<br>Guinea Savanna |  | Sudano-Sahelian |  | Tropical<br>Rainforest |  | P-value |
| --- | --- | --- | --- | --- | --- | --- | --- |
| Traits | Male | Female | Male | Female | Male | Female |  |
| Body weight | 1.48±0.05<sup>a</sup> | 1.50±0.03<sup>a</sup> | 1.53±0.03<sup>a</sup> | 1.42±0.02<sup>b</sup> | 1.43±0.04<sup>a</sup> | 1.41±0.02<sup>a</sup> | 0.049 |
| Head length | 4.54±0.09 | 4.44±0.05 | 4.79±0.05 | 4.72±0.03 | 4.67±0.06 | 4.76±0.04 | 0.147 |
| Head thickness | 10.93±0.12 | 10.75±0.07 | 11.20±0.07 | 10.98±0.05 | 11.10±0.09 | 10.98±0.85 | 0.729 |
| Helmet length | 2.57±0.11<sup>a</sup> | 2.04±0.06<sup>b</sup> | 1.96±0.06<sup>a</sup> | 1.96±0.04<sup>a</sup> | 1.98±0.08<sup>a</sup> | 2.00±0.05<sup>a</sup> | 0.001 |
| Helmet width | 1.66±0.07 | 1.61±0.04 | 1.40±0.04 | 1.35±0.03 | 1.38±0.05 | 1.29±0.03 | 0.879 |
| Wattle length | 2.53±0.09<sup>a</sup> | 2.19±0.05<sup>b</sup> | 2.34±0.06<sup>a</sup> | 2.32±0.04<sup>a</sup> | 2.28±0.07<sup>a</sup> | 2.32±0.04<sup>a</sup> | 0.015 |
| Wattle width | 1.99±0.10 | 1.77±0.06 | 1.76±0.06 | 1.56±0.04 | 1.55±0.07 | 1.51±0.04 | 0.208 |
| Neck length | 14.14±0.35 | 13.92±0.20 | 11.64±0.20 | 11.52±0.13 | 11.32±0.25 | 11.15±0.15 | 0.972 |
| Neck circumference | 8.29±0.23<sup>a</sup> | 7.76±0.13<sup>b</sup> | 7.10±0.13<sup>a</sup> | 7.33±0.08<sup>a</sup> | 7.32±0.16<sup>a</sup> | 7.13±0.10<sup>a</sup> | 0.022 |
| Wing length | 18.26±0.32 | 17.50±0.18 | 15.11±0.18 | 14.60±0.12 | 14.86±0.23 | 14.60±0.14 | 0.524 |
| Wing Span | 39.28±0.50 | 38.49±0.28 | 35.89±0.29 | 35.07±0.19 | 35.22±0.36 | 34.96±0.22 | 0.562 |
| Body length | 39.40±0.88 | 37.91±0.50 | 37.11±0.51 | 35.60±0.33 | 35.90±0.64 | 35.39±0.39 | 0.558 |
| Trunk Length | 27.52±0.39 | 26.11±0.23 | 26.44±0.23 | 25.84±0.15 | 26.06±0.29 | 25.89±0.17 | 0.092 |
| Keel length | 11.07±0.18 | 11.39±0.11 | 11.20±0.11<sup>a</sup> | 10.85±0.07<sup>b</sup> | 10.96±0.13 | 10.70±0.08 | 0.023 |
| Chest circumference | 32.60±0.69 | 31.87±0.40 | 30.32±0.40 | 29.00±0.26 | 29.07±0.50 | 28.59±0.31 | 0.516 |
| Thigh length | 11.89±0.21 | 11.38±0.12 | 11.00±0.12 | 10.70±0.08 | 10.75±0.16 | 10.62±0.09 | 0.448 |
| Shank length | 7.46±0.10 | 7.31±0.06 | 7.08±0.06 | 7.01±0.04 | 7.01±0.07 | 7.00±0.04 | 0.527 |
| Shank thickness | 4.25±0.18 | 3.95±0.10 | 5.54±0.10 | 5.36±0.07 | 5.38±0.13 | 5.39±0.08 | 0.445 |
| Massiveness | 5.40±0.21<sup>a</sup> | 5.74±0.12<sup>a</sup> | 5.86±0.12<sup>a</sup> | 5.51±0.08<sup>b</sup> | 5.51±0.15<sup>a</sup> | 5.47±0.09<sup>a</sup> | 0.038 |
| Compactness | 119.09±2.8 | 122.57±1.61 | 115.44±1.63 | 112.47±1.05 | 111.92±2.04 | 110.75±1.23 | 0.231 |
| Long-leggedness | 19.02±0.39 | 19.32±0.23 | 19.31±0.23 | 19.93±0.15 | 19.75±0.29 | 20.01±0.17 | 0.681 |
| Condition index | 8.27±0.30<sup>a</sup> | 8.81±0.17<sup>a</sup> | 10.14±0.18<sup>a</sup> | 9.69±0.11<sup>b</sup> | 9.59±0.22<sup>a</sup> | 9.64±0.13<sup>a</sup> | 0.038 |

Means within rows with P <0.05 are significantly different

### Spatial Representation of Birds

Based on Wilks’ Lambda (0.326-0.663) and F statistics (41.855-143.662) (Table 6), wing length, shank thickness, massiveness, neck circumference, head thickness, condition index, long-leggedness, neck length, thigh length and wattle length were the significant (P<0.001) parameters of importance to separate birds in the Southern Guinea Savanna, Sudano-Sahelian and Tropical Rainforest zones. However, there

**Table 6.** Traits of importance in the discriminant analysis to separate birds in the three agro-ecological zones

| **Traits** | **Wilk's Lambda** | **F-value** | **P-Level** | **Tolerance** |
| --- | --- | --- | --- | --- |
| Wing length | 0.663 | 143.662 | 0.001 | 1.000 |
| Shank thickness | 0.562 | 94.269 | 0.001 | 0.990 |
| Massiveness | 0.506 | 76.328 | 0.001 | 0.764 |
| Neck circumference | 0.465 | 65.712 | 0.001 | 0.925 |
| Head thickness | 0.420 | 61.016 | 0.001 | 0.785 |
| Condition index | 0.387 | 56.756 | 0.001 | 0.131 |
| Long-leggedness | 0.362 | 52.956 | 0.001 | 0.726 |
| Neck length | 0.343 | 49.427 | 0.001 | 0.729 |
| Thigh length | 0.332 | 45.638 | 0.001 | 0.577 |
| Wattle length | 0.326 | 41.855 | 0.001 | 0.582 |

### Contributions to variation and loadings of variables on the principal components

The result of CATPCA revealed the extraction of two principal components (PCs) which explained 42.1% of the variation in the dataset (Table 8). The first PC (Eigenvalue = 8.386) explained 27.1% of the total variance and was greatly influenced by body length (0.832), body weight (0.830), compactness (0.812), massiveness (0.810), helmet length (−0.748), wattle width (0.755), chest circumference (0.741), wattle length (−0.730), helmet width (0.723), thigh length (0.713), shank length (0.642), long−leggedness (−0.616), head thickness (0.608), condition index (0.532) and neck circumference (0.391) (Figure 3). Agro-ecology (−0.751) was more associated with the second PC (Eigen value = 4.652) which accounted for 15.0% of the total variation and had its loadings for wing length (0.754), skin colour (−0.679), neck length (0.647),, head length (−0.634), wing span (0.632), eye colour (−0.504), shank thickness (−0.490), helmet colour (0.467), helmet shape (−0.419), earlobe colour (0.390), wattle size (−0.359), plumage colour (−0.254), keel length (0.246), shank colour (0.207), trunk length (0.126). Wattle shape had equal loading for PC1 and PC2 (−0.088). However, the contributions of sex of birds to both PC1 (−0.094) and PC2 (−0.079) in terms of loadings were negligible. The high Cronbach’s alpha value of 0. 954 indicates the reliability of the CATPCA.

**Fig 2.** Canonical discriminant function illustrating the distribution of the guinea fowls among the agro-ecological zones

**Fig 3.** Individual quantitative and qualitative traits loadings on the principal components

**Table 7.**
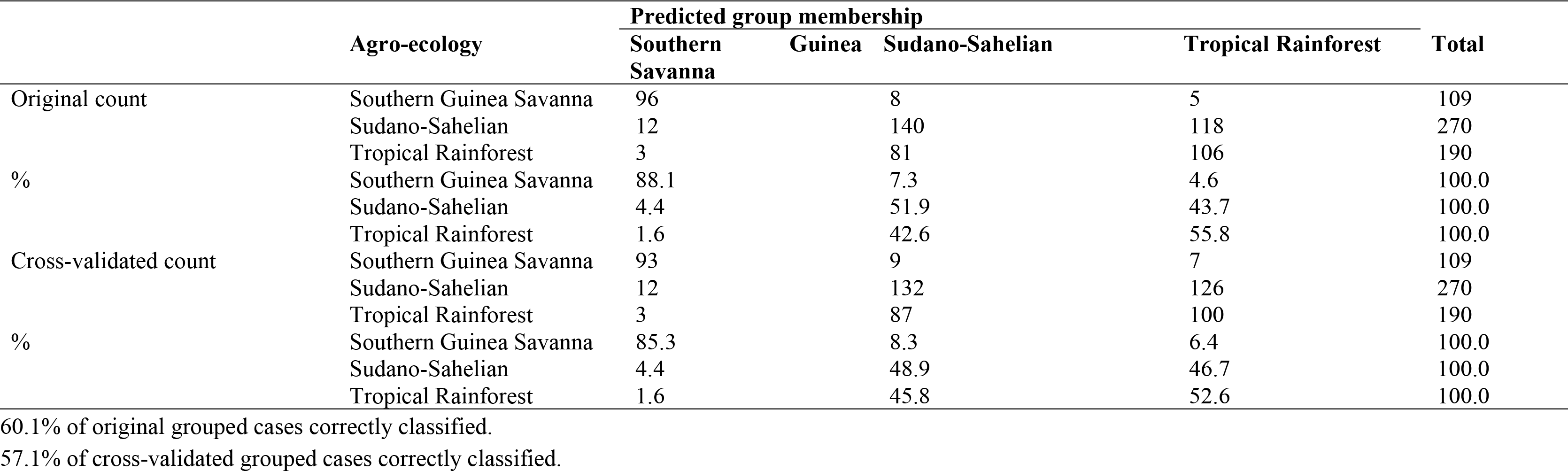
Assignment of birds to the three agro-ecological zones

**Table 8.** Eigen value and the contribution of each qualitative and quantitative trait to the total variation in the principal components

| Traits | PC1 | PC2 | Total |
| --- | --- | --- | --- |
| Plumage colour | 0.002 | 0.065 | 0.066 |
| Skin colour | 0.010 | 0.461 | 0.471 |
| Shank colour | 0.029 | 0.043 | 0.072 |
| Eye colour | 0.011 | 0.254 | 0.265 |
| Earlobe colour | 0.030 | 0.152 | 0.182 |
| Helmet colour | 0.003 | 0.218 | 0.221 |
| Helmet shape | 0.003 | 0.175 | 0.178 |
| Wattle size | 0.001 | 0.129 | 0.130 |
| Wattle shape | 0.008 | 0.008 | 0.015 |
| Body weight | 0.689 | 0.013 | 0.702 |
| Head length | 0.035 | 0.402 | 0.437 |
| Head thickness | 0.370 | 0.024 | 0.394 |
| Helmet length | 0.560 | 0.002 | 0.562 |
| Helmet width | 0.522 | 0.119 | 0.641 |
| Wattle length | 0.533 | 0.001 | 0.535 |
| Wattle width | 0.570 | 0.108 | 0.678 |
| Neck length | 0.255 | 0.419 | 0.674 |
| Neck circumference | 0.153 | 0.042 | 0.195 |
| Wing length | 0.125 | 0.569 | 0.694 |
| Wing Span | 0.053 | 0.400 | 0.452 |
| Body length | 0.692 | 0.036 | 0.728 |
| Trunk Length | 0.008 | 0.016 | 0.023 |
| Keel length | 0.028 | 0.061 | 0.089 |
| Chest circumference | 0.549 | 0.074 | 0.623 |
| Thigh length | 0.508 | 0.021 | 0.529 |
| Shank length | 0.413 | 0.016 | 0.429 |
| Shank thickness | 0.246 | 0.240 | 0.486 |
| Massiveness | 0.656 | 0.014 | 0.671 |
| Compactness | 0.660 | 0.059 | 0.719 |
| Long-leggedness | 0.380 | 0.090 | 0.470 |
| Condition index | 0.283 | 0.425 | 0.708 |
| Sex<sup>b</sup> | 0.016 | 0.561 | 0.577 |
| Agro-ecology<sup>b</sup> | 0.009 | 0.006 | 0.015 |
| Eigen value | 8.386 | 4.652 | 13.038 |
| % of Variance | 27.052 | 15.006 | 42.059 |

b= Supplementary variable.

### Decision Trees of the Data Mining

The tree diagram of the CHAID algorithm is depicted in Figure 4. Seven terminal nodes (Nodes 1, 2, 3, 5, 6, 7 and 8) were formed with the root node (Node 0) showing the descriptive statistics of the birds in the three agro-ecological zones. The Chi-squared-based branch and node distribution revealed that wing length was the variable of utmost importance in assigning the birds into their respective agro-ecological zone followed by eye colour. Wing length (>18.10 cm) only was significantly (P<0.001) sufficient to discriminate between birds of the Southern Guinea Savanna and those of Sudano-Sahelian and Tropical Rainforest zones. However, wing length (14.80-15.50 cm) together with eye colour provided a better differentiation of the Sudano-Sahelian and Tropical Rainforest zones. While birds from the former had mostly brown and pink eye colour, the later were associated mostly with white, black and bluish eye colour. With regard to model accuracy and validity, the resubstitution (probability of misclassifying an unseen instance) rate estimate of 0.420 was closely similar to the cross-validation error value of 0.431 with standard error of 0.021, respectively. However, the Exhaustive CHAID decision tree formed seven terminal nodes (Figure 5). Here, wing length (>18.10 cm) was also the best single discriminant variable (P<0.001) to distinguish birds in the three agro-ecological zones. In contrast to what was obtained under CHAID, body length and eye colour were the two additional variables to differentiate the populations. Wing length (14.80-15.50 cm), body length (<= 35.00 cm) and eye colour permitted a better separation of the Sudano-Sahelian from Tropical Rainforest birds. Unlike what was observed in CHAID, birds from the former had mostly brown, white and pink eye colour while the later were characterized by black as well as bluish eye colour. As regards model accuracy and validity, the resubstitution rate and cross-validation error estimates were 0.406 and 0.409 with standard error of 0.021, respectively.

**Fig 4.** The association between the agro-ecologies and the phenotypic traits using CHAID

**Fig 5.** The association between the agro-ecologies and the phenotypic traits using Exhaustive CHAID

## Discussion

Phenotypic variation of local animal resources indicates a genetic diversity that may be worth conserving for future uses while better understanding of the external features helps to facilitate the implementation of conservation policies aimed to ensure local resources survival [18]. Morphometric and phaneroptic approaches may be fundamental in the management of poultry considering the fact that they are fast and economically profitable [44]. The preponderance of more female birds in the present study could be attributed to the fact that smallholder poultry farmers normally keep more hens for the purpose of procreation, whereas the cocks are mostly slaughtered for consumption or sold to generate family income. We observed four major plumage colours (Pearl, Black, Brown and Pied). The varying colour patterns could be an indication that there are no pure genotypes of guinea fowl in Nigeria as there are no records of selective breeding of the indigenous stock birds. However, our colour patterns were somehow different from the dominant Pearl, Lavender, Black and White variations earlier reported in the country [48, 49]. The slight variation may be occasioned by sampling coverage. In a similar study in Ghana, Agbolosu et al. [24] found that the predominant plumage colour was pearl grey colour (43.7%), whereas Traore et al. [25] reported that pied plumage colour (42.76%) was the most frequent among the provinces in Burkina Faso. The Nigerian birds shared brown eye colour (57.0%) with those of Atakora (Mountainous) dry savannah zone in Togo [50], and black shank colour with those of Kenya (95.6%) [19], Sudanian and Sudano-Guinean zones in Benin [51].

Colour polymorphism defies evolutionary expectations as a single species may maintain a striking phenotypic variation [52]. The present variant phenotypes may be due to polymorphism [53], and might have evolved in local guinea fowls as adaptive measures for survival under varied environmental conditions. According to Getachew et al. [54] sustainable livestock production in the tropics requires adaptive genotypes which can withstand the undesirable effects of climate change and ensures optimal performance of the birds. In another study in a different species, Nigenda-Morales et al. [55] reported that the overall fitness of individuals in their environments may be affected by colour while Gong et al. et al. [56] considered colour variation as an environmental indicator, which provides clues for the study of population genetics and biogeography. The preponderance of Pearl plumage colour in our study might also be attributed to farmers’ preference. It is congruous to the submission of Vignal et al. [6] that the prevalence of a particular colour could be attached to social-cultural value without any proven relationship with a biological function. This was buttressed by the report of González Ariza et al. [44] that certain phaneroptic variables may be associated with consumers’ trends and their cultural preferences.

Our findings on helmet shape are in agreement with the report on indigenous guinea fowls in Ghanian where single shape (42.70%) predominated. The current observation on helmet shape where more females exhibited single shape is congruous to the submission of Angst et al. [57] that females have bony helmet more compact dorsoventrally while the males have taller helmet, with a more complex shape including curvature of the posterior part along the dorsoventral axis. Similarly, Agbolosu et al. [24] reported that helmet shape is more pronounced in males than females. The observation on wattle is in consonance with findings of Umosen et al. [58] that on the average females had small wattle which was mostly carried flat.

In order to ascertain the genetic purity of the birds, the MCA result did not give a perfect clustering of the birds as phenotypic homogeneity of the Guinea fowl populations was evident in Sudano-Sahelian and Tropical Rainforest birds. This is in spite of the wide geographical distance and varying environmental conditions between the two zones. This suggests that colour traits alone might not be enough to distinguish between the three agro-ecological zones. Similar submission was made by Traore et al. [25] where in spite of the enormous environmental differences; there was morphological homogeneity in qualitative traits in guinea fowls in Burkina Faso. Brown et al. [59] also observed limited phenotypic and genetic diversity in local guinea fowl in northern Ghana.

Univariate analyses revealed significant differences among zones for most zoometric traits and calculated body indices, suggesting the possible influence of these zones on the evolutionary adaptation of the sheep population in terms of these. However, there was no clear cut pattern in the linear body measurements and indices especially of the Sudano-Sahelian and Tropical Rainforest birds. The body weight values of the present study are comparable to the 1.40 kg reported by Orounladji et al. [51] for indigenous guinea fowls in a Sudanian zone in Benin, respectively. They are however, higher than the range 1.08-1.33 kg reported for adult guinea fowl (*Numida meleagris*) in a humid zone of southern Nigeria [60] and 1.275 kg obtained in Zimbabwe [61]. However, the indigenous birds are smaller in size when compared to their exotic counterparts. While Agwunobi and Ekpenyong [62] obtained a live weight of 1.5 kg in guinea fowl of ‘Golden Sovereign’ broiler strain under tropical conditions of Nigeria, Batkowska et al. [63] found a range of 2166 ± 42.5-2291 ± 46.9 kg in French commercial set. The differences may be attributed to genetics, age, physiological stage of the birds, location and management systems employed by the poultry keepers. According to Ahiagbe et al. (64), genetic make-up and management practices could affect the growth traits of guinea fowls. The exotic guinea fowls are products of many years of robust selection and breeding [65, 66]. Therefore, it is possible that crossbreeding between the indigenous and exotic will result in birds of high genetic superiority in terms of meat yield and quality, egg production and adaptation.

Sexual dimorphism provides insight into the sexual- and natural-selection pressures being experienced by male and female animals of different species [67]. At inter-population level especially with some morphometric traits, sexual dimorphism in the present study favoured male animals. This concurs with the established literature that males generally possess larger body sizes than females in normal sexual size dimorphism in birds [68]. The differential rate and duration of growth by the sexes may be responsible for the present observations. Also, high rate of breeding in the populations could be another contributing factor to sexually dimorphic traits [69] as the birds have not been selected for the purpose of classical breeding. As obtained in the current study, Dudusola et al. [60] found male dominance in thigh length, body length, wing length, wing span, wattle length and chest circumference in Nigeria while Brown et al. [59] reported longer body and shank length including wingspan in indigenous guinea fowl in Ghana. In a related study in domestic chicken, Toalombo Vargas et al. [70] reported longer thigh length in male birds.

The canonical discriminant analysis showed high level of admixture especially between the Sudano-Sahelian and Tropical Rainforest populations. It could, therefore, be reported that the guinea fowls in Nigeria are unselected and largely of mixed populations. Northern Nigeria is the traditional home of indigenous helmeted guinea fowls in the country [71]. Considering the geographical proximity of the Southern Guinea Savanna and Sudano-Sahelian zones, one would have expected considerable intermixing of the guinea fowl populations. However, the reverse was observed in the present study as the intermingling between the birds in the Sudano-Sahelian and Tropical Rainforest zones was higher which could partly be due to transhumance especially by herders. The herders (mainly cattle rearers) from the northern parts of the country do move to the southern parts in search of natural pastures during the dry season. When they do so, they tend to carry along all their animals for settlement in their new locations. In that process, there is the possibility of exchange of birds between the settlers and their hosts. Such livestock mobility, which is seen as a means to an end [72] could have shaped poultry distribution pattern.

Suffice to say that the guinea fowl (*Numida meleagris*) population of Tropical Rainforest is an ecotype of the Sudano-Sahelian; which is quite different from *Numida ptilorhycha* that is indigenous to the deciduous rain forest zone of southern Nigeria [73]. This assertion is consolidated by the reports of Ayorinde [74] and Obike et al. [75] that *Numida meleagris*, domiciled in the north was spreading to other smallholder farming areas. Yakubu et al. [30] reported that movement of herders from one State to another can impact on livestock distribution pattern in Nigeria. In a related study, Whannou et al. [76] submitted that the mobility of herders could engender genetic introgression, thereby affecting animal genetic diversity. Another possible factor that could have contributed to the genetic erosion is inter-regional trade. It appears such live animal trade seemed to be more between livestock marketers in the Tropical Rainforest and Sudano-Sahelian zones than their Southern Guinea Savanna counterparts. According to Benton et al. [77], market dynamics in one location could drive biodiversity-damaging practices in other locations. In another study, Valerio et al. [78] highlighted the relevance of cross-border ties suggesting that markets play distinct structural roles in understanding animal movement patterns.

In spite of the admixture, the results of CATPCA, CHAID and Exhaustive CHAID revealed that some levels of separation can be obtained based on agro-ecology. Wing length was identified by the three models as contributing considerably to the discrimination of the guinea fowls. However, the guinea fowls from the three agro-ecologies could best be separated using wing length, body length and eye colour. Both wing and body lengths are skeletal parameters that are not influenced by body condition, thereby providing good estimates of overall body size of the birds. It is possible that both traits are under similar selection pressure [79]. The importance of morphometric traits in population stratification has also been stressed in other avian species [80; 81].

## Conclusion

The quantitative and qualitative traits of Nigerian guinea fowls predominantly were affected by agro-ecology. However, there was no clear cut variation and distribution pattern across the three agro-ecological zones. Although birds in the Southern Guinea Savanna zone appeared to have edge over others, the indigenous birds, generally were of small body weights and morphometric traits. This could be part of the animals’ adaptation for survival under the low-inputs tropical environment. The clustering pattern of the traits especially between the Sudano-Sahelian and Tropical Rainforest birds revealed high level of admixture. This calls for further genomic studies to unravel the degree of genetic erosion and pave way for policy decisions geared towards effective management, conservation and genetic improvement of the indigenous birds. The anticipated benefits include the development of hybrid improved guinea fowls for the empowerment of women and youth including improvement in food security and livelihoods.

## Author Contributions

Conceptualization: Abdulmojeed Yakubu, Praise Jegede, Ayoola Shoyombo and Ayotunde Adebambo; Data curation: Praise Jegede, Mathew Wheto, Samuel Vincent and Harirat Mundi; Formal analysis, Abdulmojeed Yakubu and Praise Jegede; Funding acquisition: Abdulmojeed Yakubu, Mathew Wheto, Ayoola Shoyombo, Ayotunde Adebambo, Mustapha Popoola, Osamede Osaiyuwu, Olurotimi Olafadehan, Olayinka Alabi, Comfort Ukim, Adeniyi Olayanju and Olufunmilayo Adebambo; Methodology: Abdulmojeed Yakubu, Ayoola Shoyombo, Ayotunde Adebambo, Mustapha Popoola, Osamede Osaiyuwu, Olurotimi Olafadehan, Olayinka Alabi, Comfort Ukim, Osaiyuwu, Olurotimi Olafadehan, Olayinka Alabi, Comfort Ukim, Samuel Vincent, Harirat Mundi, Adeniyi Olayanju and Olufunmilayo Adebambo.

## Data Availability Statement

Data have been provided and can be found as supplementary materials

## Acknowledgments

The authors are extremely grateful to the poultry keepers, extension agents and village heads and contact persons that facilitated data collection.

## Supporting information

S1-Fig 1-5 (ZIP)

S2- Data 1-3 (ZIP)

